# Shared and reinforcer-specific alterations in the CRH and noradrenergic systems following short- and long-term withdrawal from cocaine, heroin, and sucrose self-administration

**DOI:** 10.1101/2025.10.17.683095

**Authors:** David Roura-Martínez, Marcos Ucha, Mario Moreno-Fernández, Carlos Alberto Castillo, Inmaculada Ballesteros-Yánez, Alberto Marcos, Emilio Ambrosio, Alejandro Higuera-Matas

## Abstract

Stress is known to play a critical role in relapse to drug use as well as in food craving. Craving itself is a key determinant of relapse, and cue-induced drug craving has been shown to increase, or ‘incubate’, over time for certain drugs such as cocaine and nicotine, though this effect is less consistent for others such as opiates. However, the modulations of stress-related biochemical systems after early or protracted withdrawal that could contribute to this incubation phenomenon have not yet been systematically examined in animal models, nor has the specificity of these mechanisms been tested across different drug classes or reinforcers. To address this gap, we analysed brains from male Lewis rats that self-administered cocaine (0.75 mg/kg, i.v.), heroin (0.075 mg/kg, i.v.), or saline, and subsequently assessed changes in plasma corticosterone, ornithine and other stress-related amines, alongside central gene and protein expression CRH, CRH2 receptor, and α- and β-adrenergic receptor subunits -*Adra1, Adra2a* and *Adrb1*) in cortico-striato-amygdalar nodes (after 1 or 30 days of withdrawal). A parallel experiment was conducted using sucrose as a reinforcer. Our findings indicate that although most effects were reinforcer-specific, convergent adaptations were also observed, particularly within noradrenergic systems and the basolateral amygdala, expanding our knowledge about the neurochemical rearrangements occurring during withdrawal.

## 1. INTRODUCTION

Addiction, or substance use disorder according to the DSM-5-TR terminology, has been frequently defined as a chronic relapsing disorder (Blithikioti et al., 2025; Guerrin et al., 2025; Leukefeld and Tims, 1989). As such, it is a crucial research goal to obtain a deep and comprehensive understanding of the mechanisms involved in relapse. Among other factors, stress plays a crucial role in relapse into drug use (Carmody, 1990; LaFond et al., 2024; Sinha, 2024). There is ample preclinical literature which indicates that rodents will reinstate their drug seeking behaviour after exposure to different stressors (Mantsch et al., 2016). Moreover, a previous stressful experience can facilitate reinstatement of drug-seeking behaviour (Morales-Silva et al., 2024). Indeed, seminal work demonstrated that intermittent footshock reinstates heroin seeking after extinction and that central administration of corticotropin-releasing factor (CRF) is sufficient to trigger reinstatement, whereas CRF receptor antagonism blocks stress-induced relapse (Y Shaham et al., 1997). Importantly, neither adrenalectomy nor inhibition of corticosterone synthesis blocked reinstatement, indicating that central CRF signalling, rather than stress-induced elevations in peripheral glucocorticoids, is the primary driver of stress-triggered relapse in this model. Similar findings were subsequently reported in cocaine-trained rats. Intracerebroventricular administration of a CRF receptor antagonist blocked footshock-induced reinstatement, whereas cocaine-primed reinstatement was largely unaffected, dissociating stress-triggered from drug-induced relapse mechanisms. Although adrenalectomy abolished stress-induced reinstatement, basal corticosterone replacement restored this effect, suggesting that basal glucocorticoid tone may be necessary for stress-induced reinstatement, whereas stress-induced corticosterone elevations per se are not required (Erb et al., 1998). Pharmacological specificity was subsequently strengthened by the finding that systemic administration of the selective non-peptide CRF1 antagonist CP-154,526 attenuated stress-induced reinstatement of both heroin and cocaine seeking, without affecting operant responding in the absence of stress (Y Shaham et al., 1998).

Beyond CRF signalling, converging evidence also implicated the noradrenergic system as a critical mediator of stress-induced reinstatement. In heroin self-administration -trained rats, systemic or intracerebroventricular administration of the α2-adrenergic receptor agonist clonidine, which reduces noradrenaline (NA) release, markedly attenuated footshock-induced reinstatement, whereas manipulations within the locus coeruleus were ineffective. Moreover, lesions of ventral noradrenergic projections significantly reduced stress-induced reinstatement, highlighting the contribution of extracoerulear noradrenergic pathways (Y Shaham et al., 2000). Similarly, in cocaine-trained rats, α2-adrenergic agonists such as clonidine, lofexidine, and guanabenz suppressed footshock-induced reinstatement without affecting cocaine-primed reinstatement (S Erb et al., 2000).

In humans, early clinical reports showed that stress facilitates relapse (Carmody, 1990; Wallace, 1989). The mechanisms mediating these effects are still poorly understood. Despite the preclinical evidence implicating CRF1 signalling in stress-induced reinstatement, translation to humans has proven unexpectedly challenging. In treatment-seeking alcohol-dependent patients, the CRF1 antagonist pexacerfont failed to reduce stress- or cue-induced alcohol craving, subjective distress, hypothalamic-pituitary-adrenal (HPA)-axis responses, or neural reactivity to aversive stimuli (Kwako et al., 2015). Similarly, although the CRF1 receptor antagonist verucerfont effectively suppressed HPA-axis activation and attenuated amygdala responses to negative emotional stimuli, it did not reduce stress-induced alcohol craving or behavioural markers of distress (Schwandt et al., 2016a).

In contrast to the limited clinical efficacy of CRF1 antagonists, pharmacological modulation of the NA system has yielded more consistent translational results. In opioid-dependent individuals stabilized on naltrexone, continued treatment with the α2-adrenergic agonist lofexidine reduced stress-and cue-induced craving, attenuated physiological stress reactivity, and was associated with improved abstinence outcomes (Sinha et al., 2007b). Similarly, in non-treatment-seeking cocaine users, clonidine significantly reduced stress-induced craving, with higher doses also attenuating cue-induced responses (Jobes et al., 2011a). In a larger randomized controlled trial, clonidine maintenance prolonged opioid abstinence and delayed lapse, and ecological momentary assessment analyses suggested a “decoupling” of daily-life stress from craving (Kowalczyk et al., 2015a).

An important factor driving relapse is indeed craving (Vafaie and Kober, 2022) which is a time-dependent phenomenon. Certainly, while basal craving diminishes over time, cue-induced craving increases (‘incubates’) over withdrawal time in cocaine users (Gawin and Kleber, 1986). Importantly, this phenomenon seems to be absent in opioid users (Bergeria et al., 2024) which underscores the importance of comparing across drug classes in incubation studies. While a tremendous effort has been made to search for time-dependent neurobiological mechanisms of this incubation phenomenon (Chow et al., 2025) with potential targets for treatment being identified (Liu et al., 2023), the role of stress in this paradigm and its associated biochemical mechanisms, such as the noradrenaline (NA) system and HPA axis, are still poorly understood (Venniro et al., 2021). Although modulation of stress-related systems, including CRF and noradrenergic signalling, has been shown to influence acute stress-induced relapse (see above), their contribution to the progressive, withdrawal-dependent amplification of cue reactivity that characterizes incubation of craving remains unclear. Some studies have shed new light on the mechanisms involved in incubation of seeking induced by the cessation of drug self-administration (oxycodone in these studies) following the introduction of a stressful event (an electrified barrier that punished the rats when they approached the operanda in the boxes) (Fredriksson et al., 2023; Negishi et al., 2025, 2024) but the specific role of stress-related systems was not evaluated. To address this gap, we assessed CRH and NA related markers within key cortico-striato-amygdalar nodes implicated in the incubation of craving, including the dorsomedial and ventromedial prefrontal cortex (dmPFC, vmPFC), nucleus accumbens core and shell (NAcc core, NAcc shell), and the basolateral and central nuclei of the amygdala (BLA, CeA), regions critically involved in cue-driven motivational processing and stress-related modulation of drug seeking (Chow et al., 2025). In addition, we studied the correlations among the molecular targets studied across the regions that were analysed, as we have done in previous works (Roura-Martínez et al., 2020). Moreover, we also measured peripheral mediators such as the adrenal gland index, plasma corticosterone and different amino acids relevant to metabolic processes involved in the stress response. A major strength of this work is the study of different drug types and reinforcers. Thus, in addition to comparing a psychostimulant drug (cocaine) with an opioid drug (heroin), we also introduced a third rewarding substance, sucrose-sweetened water. An incubation phenomenon similar to the one observed for cocaine has also been described for sucrose-sweetened water (Grimm, 2020; Grimm et al., 2002) and human studies have also reported incubation of food craving (Coutinho et al., 2018; Ruiz et al., 2024) which underscores the importance of including this third reward as a comparator and also due to its own significance.

Therefore, in the present study we have used a rat self-administration model previously shown to generate incubation of seeking, to study early, late, and persistent stress-related alterations across cocaine, heroin, and sucrose withdrawal. The effects observed were largely substance-specific but converged on *Adra2a* gene in the nucleus accumbens core and *Adrb1* in the basolateral amygdala (BLA). This is the first report focusing on the CRF and NA systems in withdrawal from different drug and reinforcer types, addressing a significant gap in the literature.

## 2. EXPERIMENTAL PROCEDURES

### Animals

We used 84 Lewis male rats (300-320 g at the beginning of the experiments) purchased from Harlan International Ibérica. The self-administration behaviour of these animals has been described in a previous publication (see Roura-Martínez et al., (2020b) for more details) displaying patterns of drug or sucrose intake previously shown to produce incubation of seeking. Indeed, there were no significant differences between the self-administration curves displayed by rats from both experiments (i.e. the one aimed at showing the incubation phenomenon and the one used to obtain brain samples) (p>0.35)

### Experimental design

One batch of rats was submitted to jugular catheter surgery (for drug or saline self-administration) and the other left intact (sucrose/water self-administration). After ten days of drug/saline or sucrose/water self-administration, the rats underwent 1 or 30 days of forced withdrawal with regular handling. The rats that received intravenous administration of cocaine, heroin, or saline (iv control) were segregated into six groups (3 substances × 2 withdrawal periods; n=8 rats per group), while the rats that received sucrose or water (oral control) were distributed in four groups (2 substances × 2 withdrawal periods; n=9 rats per group).

### Surgical procedure

An intravenous polyvinylchloride tubing (0.064 mm i.d.) catheter was implanted into the right jugular vein at approximately the level of the atrium and passed subcutaneously to exit the midscapular region (Higuera-Matas et al., 2008). Surgical procedures were performed under isoflurane gas anaesthesia (5% for induction and 2% for maintenance) and buprenorphine analgesia, and the catheters were flushed daily with 0.5 mL of gentamicin (40 mg/mL) dissolved in heparinized saline in order to prevent infection and to maintain patency (see (Roura-Martínez et al., 2020)).

### Self-administration

Self-administration sessions were performed in Skinner boxes (Coulbourn Instruments or Med-Associates) equipped with two levers (active and inactive) monitored with Med-PC software. The house light was off during the sessions, and the door of the sound-attenuating cubicle left ajar, allowing some environmental light from the room. Each time the active lever was pressed (fixed-ratio 1), a pump outside the box was switched on for 5 seconds and either the drug or saline solution was infused through the catheter, or the sucrose solution or water was dispensed into a receptacle placed in between the levers. A cue-light over the active lever switched on for 10 seconds at the same time. Subsequently, there was a time-out period of 40 seconds with no programmed consequences. Cocaine, heroin, or saline self-administration sessions lasted 6 hours per day, as described previously. (Roura-Martínez et al., 2020). Rats orally self-administering sucrose or water were subjected to shorter sessions (2 h/d). The doses per injection used in the experiments were 0.075 mg/kg of heroin; 0.75 mg/kg of cocaine-HCl; and 10% w/v sucrose (Sigma-Aldrich S1888), diluted in 0.1 mL of saline (0.9%, NaCl physiological saline). To facilitate the acquisition of this behaviour, we placed two sucrose pellets on the active lever in the first two self-administration sessions.

### Animal sacrifice and tissue processing

We weighed and sacrificed the rats after 1 or 30 days of withdrawal by decapitation, between 11:00 and 13:00. Immediately after, the brain was removed in about one minute, frozen by ten seconds of immersion in isopentane (VWR 24872.298) cooled with dry ice, and stored at -70°C. We collected trunk blood in a heparinized tube and extracted plasma by centrifugation at 1000g for 10 min at 4 °C, then stored at -70°C. Adrenal glands were removed and weighed.

### Brain dissection

Brain was equilibrated 1 h at -20°C in the cryostat chamber (Microm, Cryostat HM 500O) embedded in TissueTek (Sakura, 4583). We collected 300μm thickness slices on a sterile cold surface and dissected them with a lancet, according to a rat neuroanatomical atlas (Paxinos and Watson, 2007; regions depicted in Figure 6). We stored the tissue during dissection in sterile tubes of 0.2 mL in dry ice and then at -70°C.

### Tissue homogenization

We weighed and homogenized the samples in HEPES buffer (50 mM, pH 7.5) prepared in DEPC-treated water, containing sucrose (320 mM), protease and phosphatase inhibitors, and sodium butyrate (20 mM) using a motorized mortar (Sigma-Aldrich, pellet pestle Z359971). After being kept on ice for 10 min they were centrifuged at 1000 g for 10 min at 4°C. A volume of supernatant equivalent to 3-4 mg of original tissue (≤80 μL) was added to 800 μL of QIAzol (Qiagen 79306) for extraction of cytoplasmic RNA. Another volume was added to loading buffer (Tris 62.5 mM pH 6.8, glycerol 20% v/v, SDS 2% w/v, dithiothreitol 50 mM) for Western blot. Then, we stored the samples at -70°C.

### Level of plasma corticosterone by radioimmunoassay

The plasmas were thawed, centrifuged at 10,000 g for 10 min at 4°C to discard precipitates, allowed to reach room temperature (RT), and then analysed using the Coat-a-Count® Rat Corticosterone kit (Siemens). Briefly, 50μL of sample or calibrator was added to tubes covered with anti-corticosterone antibody. Then, 1 mL of I^125^-corticosterone solution was added, vortexed and incubated at RT for 2 h. The content was then decanted and the signal analysed (1 min/tube, in counts per minute, cpm) in a gamma counter (Wallac, Automatic Gamma Counter 1470 Wizard). To estimate the non-specific binding, samples or calibrators in non-covered tubes were measured. The maximum binding value was estimated by measuring a calibrated tube without labelled corticosterone. The corticosterone values were interpolated in a semi-logarithmic line plotted against the logarithm of the concentration with the binding value and expressed in ng/mL.

### Level of plasma amines by capillary electrophoresis

The content of amines was analysed according to the method described in Lorenzo et al. (2013) (Lorenzo et al., 2013). Samples were passed through a 0.22 μm filter (VWR 28145-491 13 mm syringe filter 0.2 μm PTFE membrane) and then stored at -70°C until use. In parallel, mixtures of different amines were prepared to be used as a calibration curve (from Sigma: L-glutamic acid (49449), glycine (G7126), L-glutamine (49419), taurine (T0625), L-serine (S4500), D-serine (S4250), L-proline (81709), L-isoleucine (W527602), L-ornithine (O2375), L-threonine (T8625)). L-2-aminoadipic acid (Sigma, A7275) was used as internal standard (IS). In each cocktail, the concentration of each amine was different, but the total concentration of amines was maintained constant. The derivatization (20 μL sample, 20 μL 200 mM IS, 25 μL 40 mM 4-fluoro-7-nitro-benzofurazan (Alfa-Aesar J61336), 150 μL 10 mM borate buffer pH 10.0) was carried out at 60°C for 15 min in darkness. Then, the samples were stored in the chamber of the capillary electrophoresis apparatus (Beckman Coulter PA 800 plus) at 7°C for at least 30 min before the start of the electrophoresis, carried out at 17°C (electrophoresis buffer: 175 mM borate buffer pH 10.25, 12.5 mM β-cyclodextrin (SAFC W40,282-6)). Before its first use, the capillary column of silica (length: 60cm; inner diameter: 75μm) was conditioned with 1 M NaOH (15 min) and water (15 min), and between electrophoresis with 0.1 M HCl (3 min), water (5 min) and electrophoresis buffer (5 min). The sample was injected at the anode at 0.5 psi (33 mbar) for 10 seconds. A potential difference of 21 kV was applied, observing currents of 120 μA. The molecules were detected at the cathode by means of laser induced fluorescence (LIF), exciting at 488 nm (argon lamp) and detecting the emission at 522 nm. To obtain reproducible electropherograms, the electrophoresis buffer was renewed every six analyses. In each session, a calibration curve was also injected. The electropherograms were analysed with the 32 KaratTM software, using the corrected area normalized to the internal standard. The values were expressed as pmol/mL.

### Isolation of RNA

The RNA was extracted using a protocol adapted from Chomczynski and Sacchi (Chomczynski and Sacchi, 2006, 1987). Briefly, the sample (stored in QIAzol) was thawed and left at RT for at least 5 min. 160 μL of chloroform (Merck 1.02445.2500) were added, mixed for 15 seconds and incubated at RT for 2-3 min. It was then centrifuged at 12,000 g for 15 min at 4°C. The upper (aqueous) phase was transferred to a new sterile 1.5 mL tube, with 400 μL of isopropanol (Fischer Scientific, BP2618) and 10 μg of glycogen (Sigma G1767), mixed and incubated at RT for 10 min. It was centrifuged at 12000 g for 10 min at 4°C and the supernatant discarded. The precipitate was washed twice in 1 mL of cold (-20°C) 75% v/v ethanol and centrifuged at 7500g for 5 min at 4°C, discarding the supernatants. Finally, the precipitate was allowed to air-dry, resuspended in nuclease-free water and stored at -70°C. The RNA was quantified in a Bioanalizer 2100 system (Agilent, RNA Nano Chips, 5067-1511), with values of RIN≥7.0. Unless exceptions, absorbance ratios 260nm/280nm ≥1.8 were obtained. An isolation efficiency of 0.7-1.0 μg RNA/mg tissue was obtained.

### RT-qPCR

About 250-500ng RNA (depending on the region) was treated with DNaseI (Invitrogen, 18068-015), retrotranscribed (Biorad, iScript cDNA Synthesis kit, 170-8891), diluted 1:10 in nuclease-free water and stored at -70°C. Gene expression was analysed using SYBR Green (Biorad, SsoAdvanced Universal SYBR Green Supermix), in a CFX96 C1000 Touch of Biorad system. The total reaction volume was 10μL, and the primers were in the range of 500-750nM. The thermal protocol was as follows: 5 min at 95°C, 40 cycles of 5 s at 95°C, and 30 s at 60°C, followed by a melting curve. We calculated the Ct values and the efficiencies with the free software LinRegPCR (Ruijter et al., 2009) and then calculated the fold change for each sample and gene according to Pfaffl (Pfaffl, 2001), using Gapdh as housekeeping gene (which did not vary across conditions). Because adrenoceptors have different (and sometimes opposite) effects in different brain regions, we also calculated ratios between receptors for each region and ratios between regions for each receptor.

#### Primer list

α1 adrenoceptor (Adra1).

S: 5’-ctcgagagaagaaagctgcca-3’ A: 5’-aaaacggtttccgaaggcttg-3’

α2A Adrenoceptor (Adra2a).

S: 5’-tcctgagagggaagggattt-3’ A: 5’-agttactggggcaagtggtg-3’

β1 adrenoceptor (Adrb1).

S: 5’-gctctggacttcggtagacg-3’ A: 5’-acttggggtcgttgtagcag-3’

### Protein quantification

Protein content was quantified in microplates using the colorimetric method described by Bradford (Bradford, 1976), with minor modifications. Briefly, 250 μL of the reagent (a 1:5 dilution of the Bio-Rad Protein Assay Dye Reagent Concentrate 500-0006) was added to 5 μL of the sample. A calibration curve of BSA was made. The 595nm/450nm absorbance ratio (quantified in an Asys Hitech DigiScan) was used to interpolate unknown values.

### Western blot

5 μg of protein from each sample was resolved on 12% SDS-PAGE gels in a MiniProtean® Tetra System (Bio-Rad). To quantify the total protein, the gels were exposed to UV light for about 1 min to activate the reaction between trichloroethanol and tryptophan (Ladner et al., 2004), and they were then recorded on an Amersham Imager 600 (General Electrics). The proteins were transferred to 0.2 μm PVDF membranes (Trans-Blot® TurboTM transfer pack, Bio-Rad) in a Trans-Blot® Turbo transfer system (Biorad), blocked with 5% w/v BSA in TBST for 1 h at RT and then probed overnight in TBST at 4 °C with the primary anti-CRH antibody (Abcam Cat# ab184238, RRID:AB_2827511) or anti-CRH receptor 2 (Abcam Cat# ab104368, RRID:AB_10973387) diluted 1:3.000. Antibody binding was recognized for one hour at RT with an HRP-conjugated anti-rabbit IgG secondary antibody (Abcam Cat# ab6721, RRID:AB_955447) diluted 1:5,000, and the signal was developed using ECL2 and recorded on an Amersham Imager 600. Proteins were analysed by densitometry using ImageJ free software and the data were normalized to total protein and by gel according to the criterion described by Degasperi (Degasperi et al., 2014).

### Missing samples

Due to storage problems, two brains were lost (water wd1 group).

### Univariate analysis

We carried out two-way ANOVA (IBM Statistics) to compare specific biochemical parameters (log or inverse transformation when necessary) between groups due to the drug exposure -drug- (cocaine, heroin or saline), sucrose exposure -sucrose- (sucrose vs water) and/or withdrawal (wd1 vs wd30). Interactions were analysed by simple effects analysis (Šídák correction). Following a significant effect of the treatment, we performed the Ryan post hoc procedure (REGWQ) to allocate the differences between groups (Field, 2013).

### Bivariate statistics

To detect correlated variables that could have a psychobiological meaning, an initial screening of Pearson correlations (IBM Statistics) followed by their respective comparisons between groups was performed. The Fisher transformation was applied to the r-value, and then the Z-value and its p-value were calculated to compare the correlations between groups. We made two types of comparisons for each substance: a) the wd1 group versus its wd1 control and its wd30 group, and b) the wd30 group versus its wd30 control and its wd1 group. To select the correlations with differences between groups, no multiple comparisons correction was applied but we filtered them on the basis of three criteria: 1) n>5 in the three compared groups; 2) a significant r (p<0.05) in the group to be tested; and 3) a significant Z (p<0.05) for comparisons.

## 3. RESULTS

For all variables, descriptive statistics (N, mean, and SD per group) and inferential results are presented in Tables 1 (saline, cocaine, heroin) and 2 (water, sucrose). Table 3 shows the parameters from the correlation analyses. For the sake of conciseness, we only report here the statistics of the most relevant interactions and post-hoc tests.

The behavioural data from these animals has been published elsewhere (Roura-Martínez et al., 2020). Importantly, rats had self-administration patterns that were indistinguishable from those of rats displaying the incubation phenomenon (i.e. more lever presses under extinction conditions in wd30 compared to wd1), for all the reinforcers tested (cocaine, heroin and sucrose) (see Roura-Martínez et al. (2020) for further details).

### 3.1. Stress-related peripheral parameters

#### 3.1.1. Body weight

Differences in body weight were only attributable to the passage of time (main effect of withdrawal), with no effects of drug or sucrose nor of their interaction with withdrawal time (Tables 1 and 2).

#### 3.1.2. Adrenal gland index

In parallel with weight gain, the adrenal gland index decreased over time. In addition, this index was higher in the cocaine and heroin groups during early withdrawal (wd1) (drug × withdrawal: F(2,42)=5.68, p=0.007, η²p=0.21; saline wd1 vs cocaine wd1, p<0.001; saline wd1 vs heroin wd1, p<0.001). We did not detect any significant differences in the sucrose experiment (Fig. 1A and B; See Tables 1 and 2 for further details).

**Figure 1.**
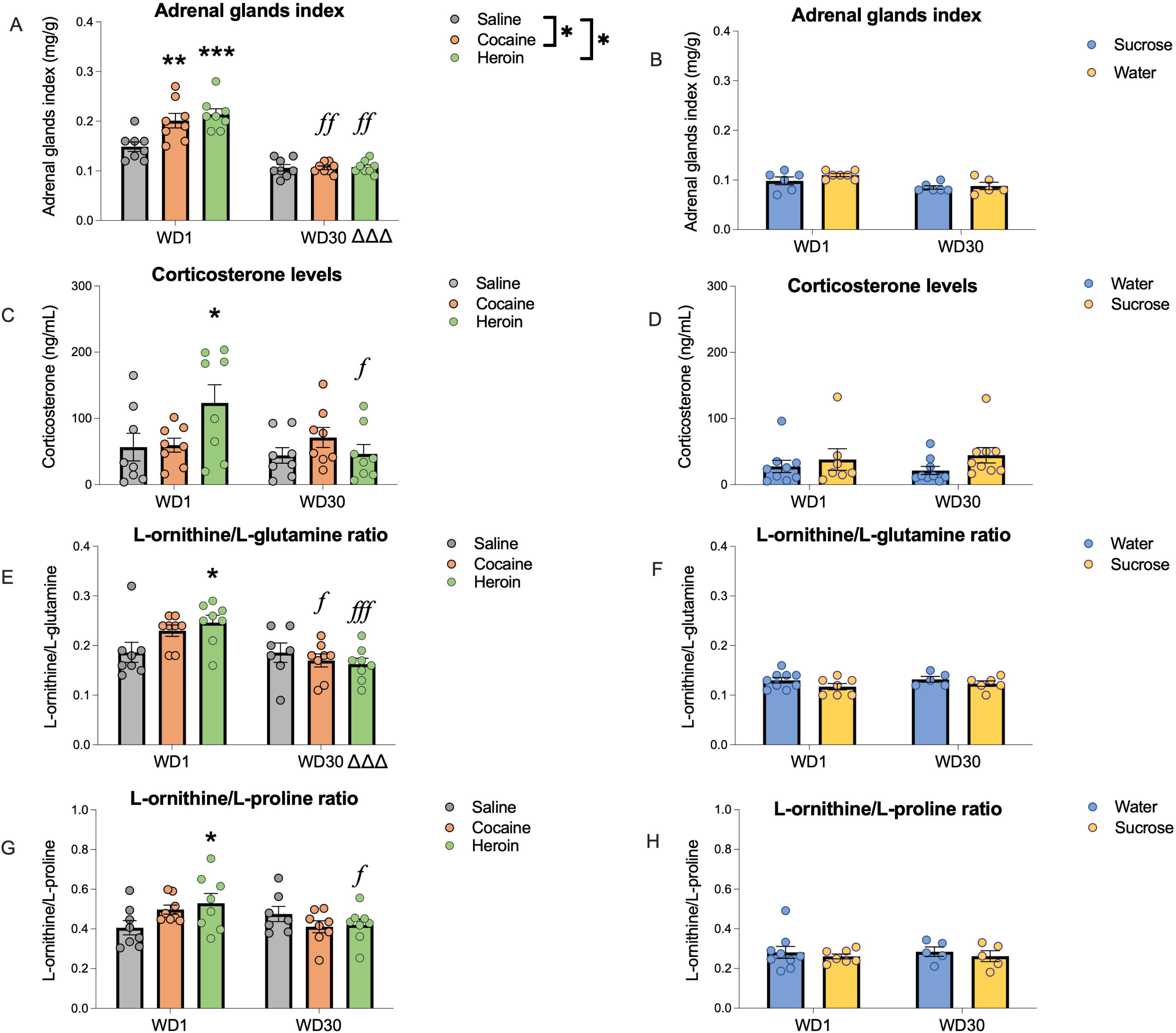
Changes in stress-related peripheral parameters during withdrawal. Levels of adrenal glands index (A, B), plasma corticosterone (C, D), and L-ornithine metabolism ratios (E-H) during cocaine and heroin (A, C, E, G) or sucrose (B, D, F, H) withdrawal. Individual values are presented as well as the mean ± SEM. Differences relative to the control group, *p<0.05, **p<0.01, ***p<0.001; differences relative to the same treatment on different withdrawal days, *f* p<0.05, *ff* p<0.01, *fff* p<0.001; general differences between the two withdrawal days, Δ p<0.05, ΔΔ p<0.01, ΔΔΔ p<0.001.

#### 3.1.3. Corticosterone

We observed a significant drug × withdrawal interaction (F(2,42)=3.32, p=0.046, η²p=0.14) which, upon further analysis, showed that plasma corticosterone levels were significantly higher in the heroin group compared with the saline group during early withdrawal (saline wd1 vs heroin wd1, p=0.033). Moreover, a significant effect of withdrawal was observed in the heroin group (Fig. 1C), with a reduction in corticosterone levels over time (heroin wd1 vs heroin wd30, p=0.004). There were no effects in the sucrose/water experiment (Fig. 1D) (Tables 1 and 2).

#### 3.1.4. Plasma amino acid levels and L-ornithine metabolism

Among the amines analysed, we found significant modulations in those related to L-ornithine metabolism. This amino acid is metabolically linked to L-glutamate, L-glutamine, and L-proline (Watford, 2008), and is also associated with stress (see Discussion). We analysed both the concentrations of these amines individually and the ratios of L-ornithine to each of the others. Cocaine, heroin, or sucrose self-administration did not alter absolute concentrations (Tables 1 and 2). However, when examining ratios, we found an increment in the L-ornithine/L-glutamine ratio for heroin at early withdrawal compared with saline and decreases across withdrawal for both drugs (significant drug × withdrawal interaction: F(2,41)=3.77, p=0.031, η²p=0.16; saline wd1 vs heroin wd1, p=0.024; cocaine wd1 vs cocaine wd30, p=0.008; heroin wd1 vs heroin wd30, p<.001) (Fig. 1E). Similarly, the L-ornithine/L-proline ratio increased at early withdrawal but decreased after one month of withdrawal in heroin rats (drug × withdrawal: F(2,41)=3.71, p=0.033, η²p=0.15; saline wd1 vs heroin wd1, p=0.048; heroin wd1 vs heroin wd30, p=0.032 Fig. 1G). No significant effects were observed in the sucrose/water experiment (Fig. 1F and 1H; Tables 1 and 2).

### 3.2. Stress-related central parameters

#### 3.2.1. Enduring self-administration effects

All three rewarding substances tested (cocaine, heroin and sucrose) produced long-lasting effects in the nucleus accumbens. In terms of gene expression, cocaine self-administration was associated with a trend towards decreased *Adra2a* expression levels in the shell (Fig. 2C; drug effect: F(2,38)=3.55, p=0.039, η²p=.16; saline vs cocaine, p=0.068) and an increase in the *Adra2a* core/shell ratio (Fig. 2G; drug effect: F(2,36)=3.65, p=0.036, η²p=.17; saline vs cocaine, p=0.033). This ratio also increased after sucrose consumption (Fig. 2H; sucrose effect: F(1,19)=8.18, p=0.010, η²p=0.30), accompanied by additional changes in the core: decreases in *Adra1* (Fig. 2B; sucrose effect: F(1,30)=9.28, p=0.005, η²p=0.24) and increases in the *Adra2a/Adra1* ratio (Fig. 2F; sucrose effect: F(1,21)=7.82, p=0.011, η²p=0.27).

**Figure 2.**
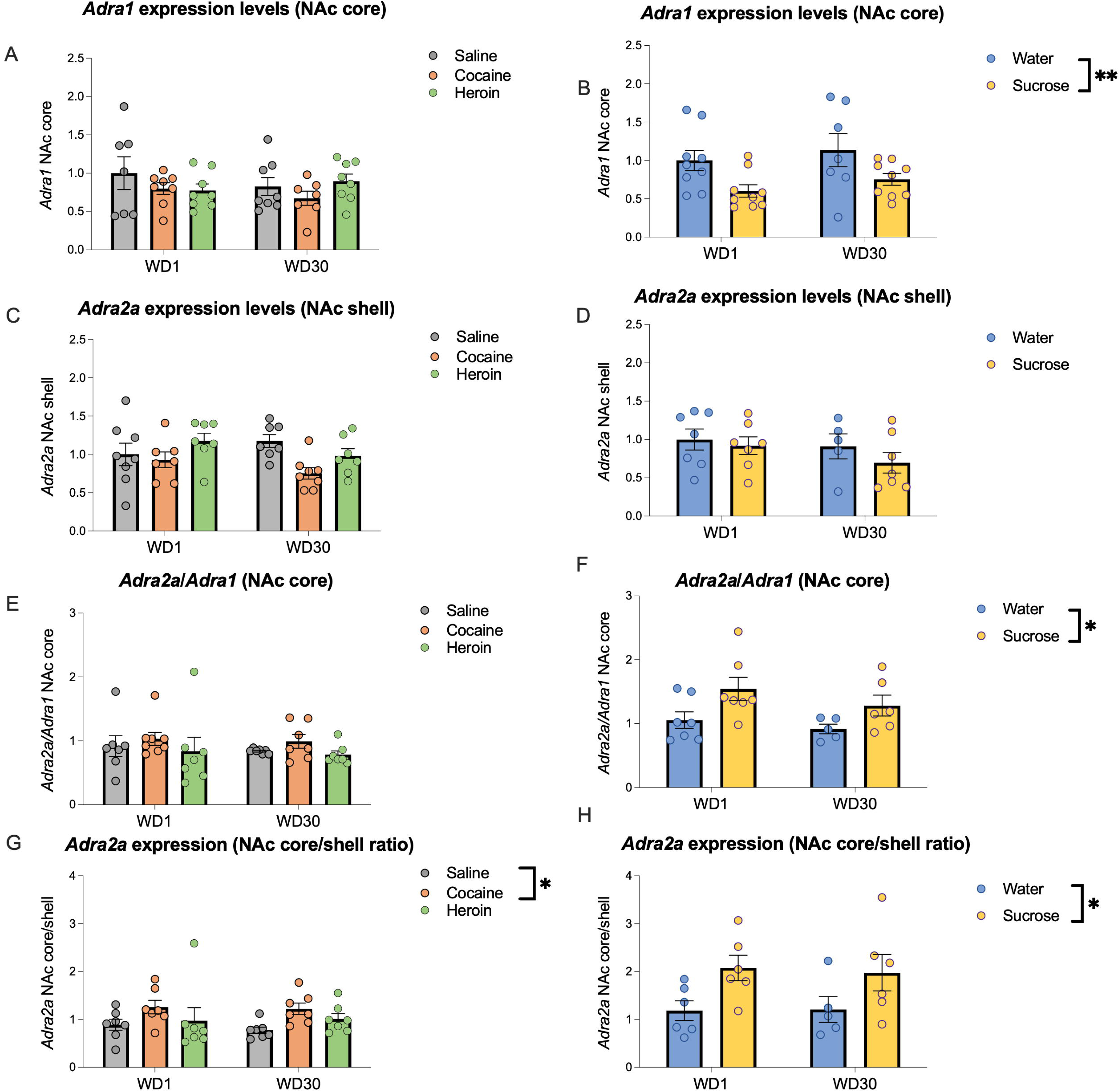
Changes in gene expression of adrenoceptors in the nucleus accumbens during withdrawal. Levels of *Adra1* NAc core (A, B), *Adra2a* NAc core (C, D), *Adra2a*/*Adra1* NAc core (E, F), and *Adra2a* NAc core/shell (G, H) during cocaine and heroin (A, C, E, G) or sucrose (B, D, F, H) withdrawal. Individual values are presented as well as the mean ± SEM. Differences relative to the control group, *p<0.05, **p<0.01, ***p<0.001.

In the prefrontal cortex, sucrose self-administration also produced lasting changes in the dmPFC: decreases in *Adrb1* (Fig. 3F; sucrose effect: F(1,30)=21.40, p<0.001, η²p=0.42), *Adra1* (Fig. 3B; sucrose effect: F(1,30)=9.96, p=0.004, η²p=0.25), *Adra2a* (Fig. 3D; sucrose effect: F(1,30)=8.48, p=0.007, η²p=0.22), *Adra1/Adrb1* ratio (Fig. 4F; sucrose effect: F(1,30)=24.37, p<.001, η²p=0.45), and *Adra1* dmPFC/vmPFC ratio (Fig. 4H; sucrose effect: F(1,30)=10.14, p=0.003, η²p=.26); and increases in the *Adra2a/Adra1* (Fig. 4B; sucrose effect: F(1,30)=29.38, p<.001, η²p=.50), *Adra2a* BLA/dmPFC ratio (Fig. 4J; sucrose effect: F(1,22)=8.94, p=0.007, η²p=0.29), and dmPFC *Adra2a*/*Adra1* ratio (Fig. 4D; sucrose effect: F(1,30)=4.53, p=0.042, η²p=0.13).

**Figure 3.**
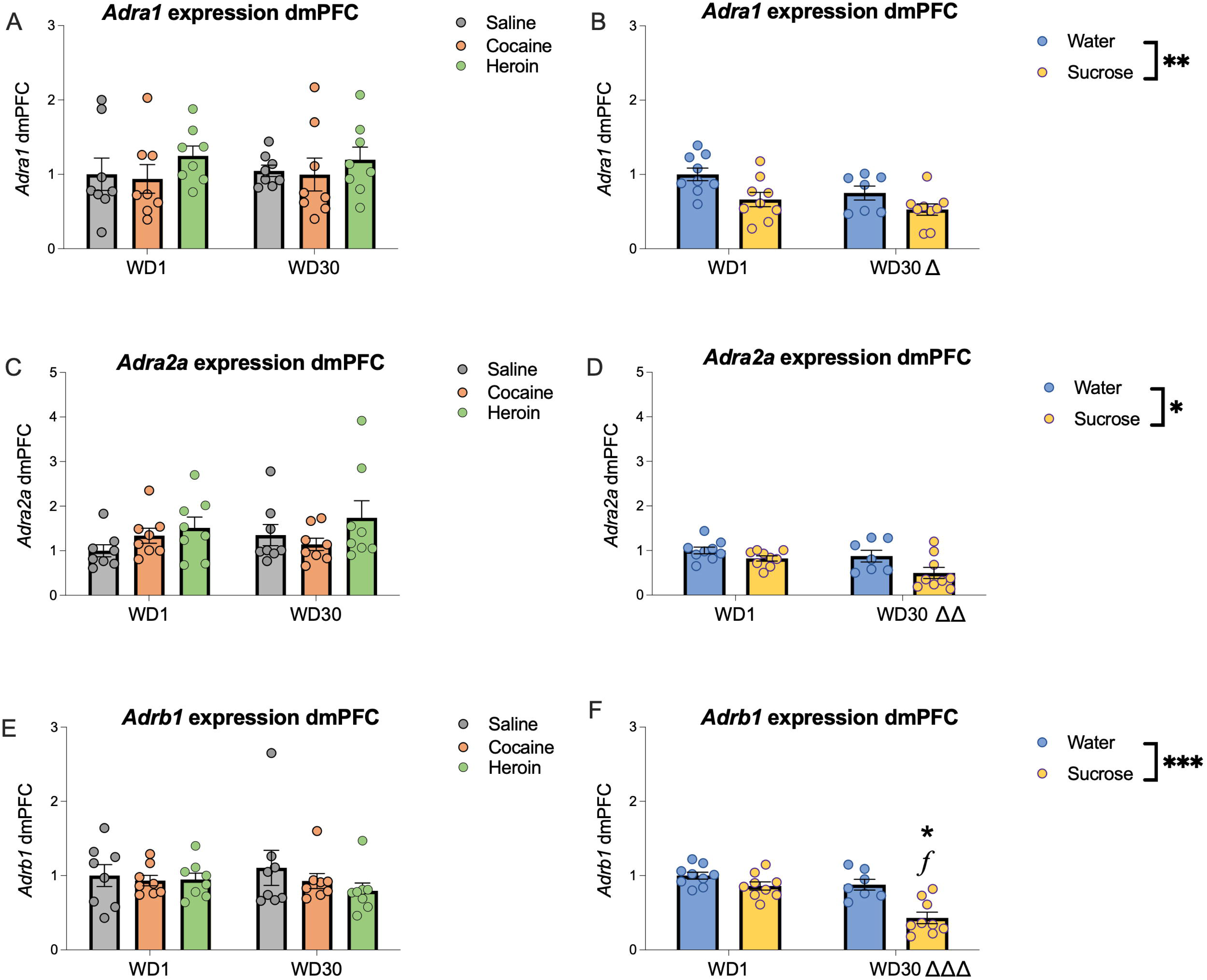
Changes in gene expression of adrenoceptors in the dorsomedial prefrontal cortex during withdrawal. Levels of *Adra1* (A, B), *Adra2a* (C, D), and *Adrb1* (E, F) during cocaine and heroin (A, C, E) or sucrose (B, D, F) withdrawal. Individual values are presented as well as the mean ± SEM. Differences relative to the control group, *p<0.05, **p<0.01, ***p<0.001; differences relative to the same treatment on different withdrawal days, *f* p<0.05, *ff* p<0.01, *fff* p<0.001; general differences between the two withdrawal days, Δ p<0.05, ΔΔ p<0.01, ΔΔΔ p<0.001.

**Figure 4.**
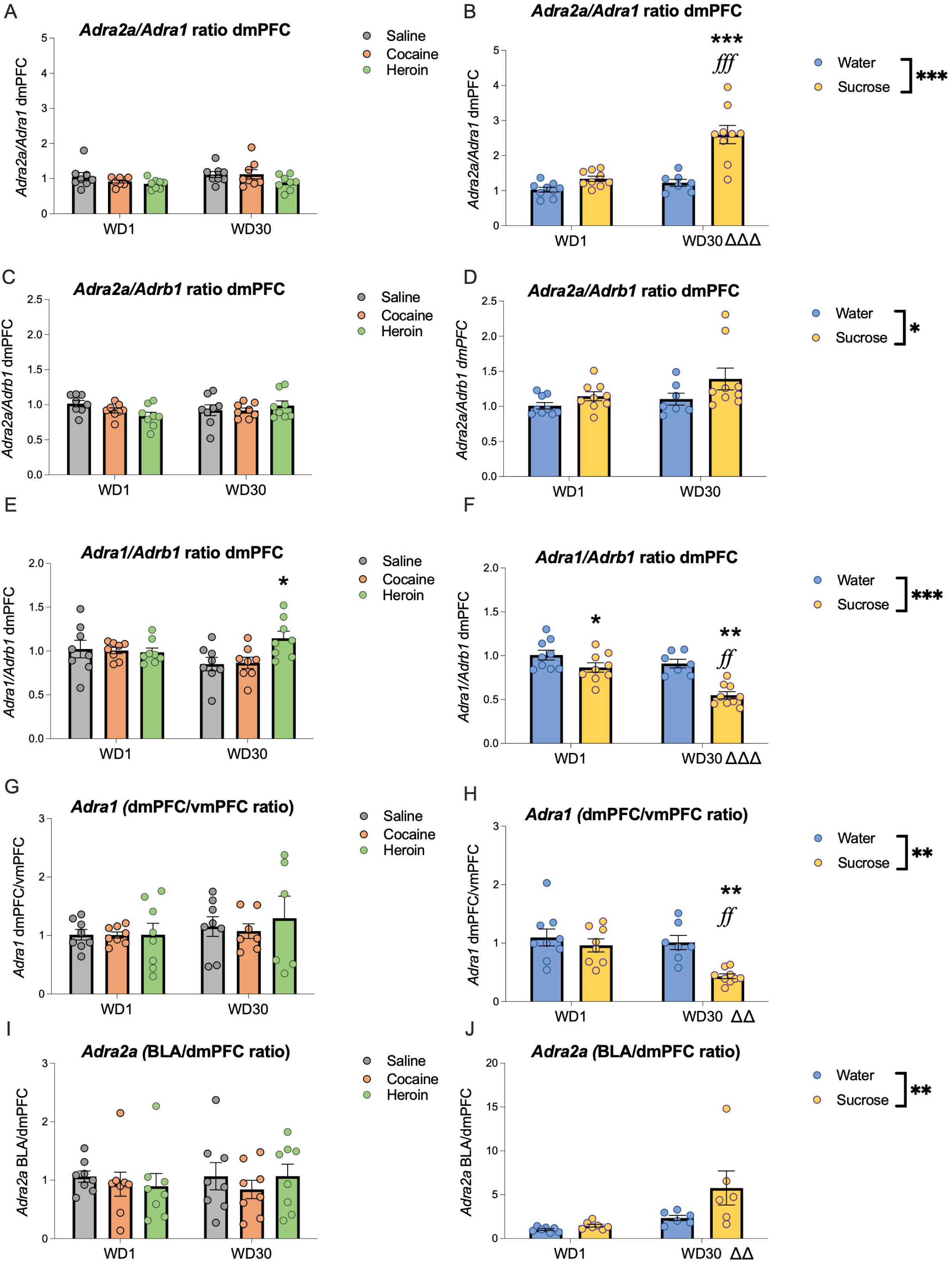
Changes in gene expression of adrenoceptors in the dorsomedial prefrontal cortex (and in relation with BLA) during withdrawal. Levels of *Adra2a*/*Adra1* dmPFC (A, B), *Adra2a*/*Adrb1* dmPFC (C, D), *Adra1*/*Adrb1* dmPFC (E, F), *Adra1* dmPFC/vmPFC (G, H), and *Adra2a* BLA/dmPFC (I, J) during cocaine and heroin (A, C, E, G, I) or sucrose (B, D, F, H, J) withdrawal. Individual values are presented as well as the mean ± SEM. Differences relative to the control group, *p<0.05, **p<0.01, ***p<0.001; differences relative to the same treatment on different withdrawal days, *f* p<0.05, *ff* p<0.01, *fff* p<0.001; general differences between the two withdrawal days, Δ p<0.05, ΔΔ p<0.01, ΔΔΔ p<0.001.

We then computed correlations between measured parameters and examined their modulation by drug/sucrose self-administration and withdrawal (Table 3). In the nucleus accumbens core, CRH and *Adra2a* levels were negatively correlated in saline groups (saline wd1, r=–0.949, p=0.001; saline wd30, r=–0.779, p=0.023). This correlation was abolished following cocaine (as evidenced by the significant difference among the correlations revealed by the differences among z scores) (wd1: Z=–2.533, p=0.011; wd30: Z=–3.079, p=0.002), and also by heroin self-administration (wd1: Z=–2.411, p=0.016; wd30: Z=–1.983, p=0.047) (Fig. 5).

**Figure 5.**
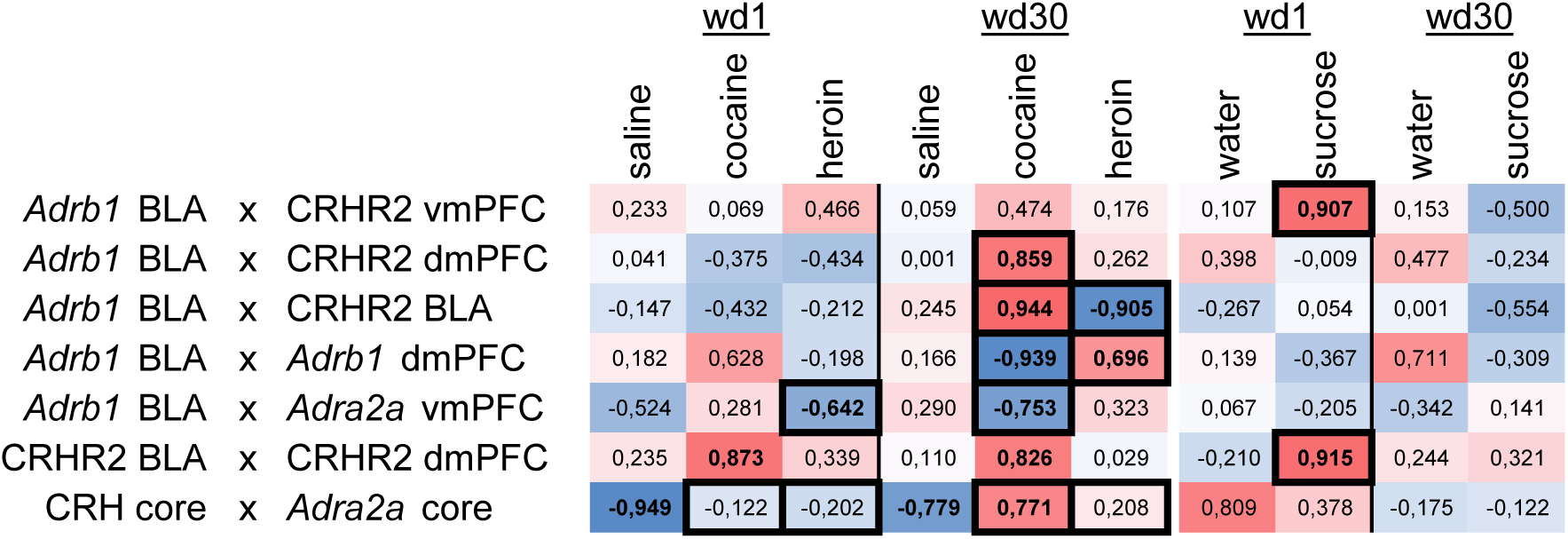
Heatmap of correlations between different parameters. For each pair of variables that experienced changes, we report the r value of the correlation for each experimental group. A bold value indicates a significant r value; a boxed value indicates a significant change relative to their control groups (control treatment and abstinence day).

#### 3.2.2. Modulations occurring across withdrawal

Specific dmPFC alterations that were emerging after sucrose self-administration, albeit non-significantly, became significant after one month of withdrawal. These changes included decreases in *Adrb1* gene expression (Fig. 3F; sucrose × withdrawal: F(1,30)=5.93, p=0.021, η²p=0.17; water wd30 vs sucrose wd30, p<0.001; sucrose wd1 vs sucrose wd30, p<0.001), a trend for a decreases in the *Adra1* dmPFC/vmPFC ratio (Fig. 4H; sucrose × withdrawal: F(1,29)=3.90, p=0.058, η²p=0.12; water wd30 vs sucrose wd30, p=0.001; sucrose wd1 vs sucrose wd30, p=0.002), and reductions in the *Adra1*/*Adrb1* ratio (Fig. 4F; sucrose × withdrawal: F(1,30)=4.68, p=0.039, η²p=0.14; trend in water wd1 vs sucrose wd1, p=0.051; water wd30 vs sucrose wd30, p<0.001; sucrose wd1 vs sucrose wd30, p<0.001); In addition, there were also increments in the *Adra2a*/*Adra1* ratio (Fig. 4B; sucrose × withdrawal: F(1,30)=11.69, p=0.002, η²p=0.28; water wd30 vs sucrose wd30, p<0.001; sucrose wd1 vs sucrose wd30, p<0.001). Moreover, the *Adra1*/*Adrb1* ratio also increased after one month of heroin withdrawal (Fig. 4E; drug × withdrawal F(2,42)=3.34, p=0.045, η²p=0.14; saline wd30 vs heroin wd30, p=0.016).

When we analysed the patterns of correlations across withdrawal, we observed, in the nucleus accumbens core, a significant correlation between CRH and *Adra2a*, specifically in the cocaine wd30 group (r=0.77,1 p=0.042). Most importantly, we found a common node of changes in correlations across all reinforcers, centred on *Adrb1* expression in the BLA. In addition, we found that sucrose self-administration and withdrawal modified the correlations between *Adrb1* in BLA and CRHR2 in the vmPFC, as evidenced by the significant difference between z scores (water wd1 =0.107, p=0.819; sucrose wd1, r=0.907, p=0.005; wd1, water vs sucrose, Z=–1.983, p=0.047; sucrose wd1 vs sucrose wd30, Z=2.697, p=0.007), and that cocaine also modified the correlations between BLA *Adrb1* expression levels and CRHR2 accumulation in the dmPFC (saline wd30 r=0.001, p=0.998; cocaine wd30, r=0.859, p=0.006; wd30, saline vs cocaine, Z=–2.038, p=0.042; cocaine wd1 vs wd30, Z=–2.662, p=0.008). In addition, the correlation between BLA *Adrb1* and CRHR2 levels changed during cocaine withdrawal (cocaine wd30, r=0.944, p<0.001; wd30, saline vs cocaine, Z=–2.270, p=0.023; cocaine wd1 vs wd30, Z=–3.060, p=0.002), as was the case also with heroin, which modified correlations in the BLA (heroin wd30, r=–0.905, p=0.005; wd30, saline vs heroin, Z=2.471, p=0.013 and a trend in heroin wd1 vs wd30, Z=1.910, p=0.056). BLA *Adrb1* correlations with *Adrb1* in the dmPFC and with *Adra2a* in the vmPFC also shifted during withdrawal from both drugs of abuse (Fig. 5; Detailed statistics are provided in Table 3).

## 4. DISCUSSION

The main aim of this study was to characterise the modulations in CRF and NA systems occurring during withdrawal from cocaine, heroin and the natural reinforcer sucrose, which may be relevant to the incubation of seeking phenomenon. Using a rat model of self-administration which generated intake patterns previously found to produce incubation of seeking (Roura-Martínez et al., 2020), we identified changes emerging both at early and late stages of withdrawal, as well as persistent effects. Some of these alterations were shared between two of the reinforcers tested, but most were specific. For example, during early drug withdrawal (cocaine and heroin), we observed adrenal hypertrophy and disruptions in ornithine metabolism. Nevertheless, common nodes emerged across all reinforcers: ***Adra2a*** expression in the nucleus accumbens core (long-lasting effects) and ***Adrb1*** expression in the BLA (late effects) (Fig. 6). This study addresses a gap in the literature, as few investigations on incubation of seeking have examined peripheral or central stress-related parameters, despite the well-established role of stress in modulating drug seeking.

**Figure 6.**
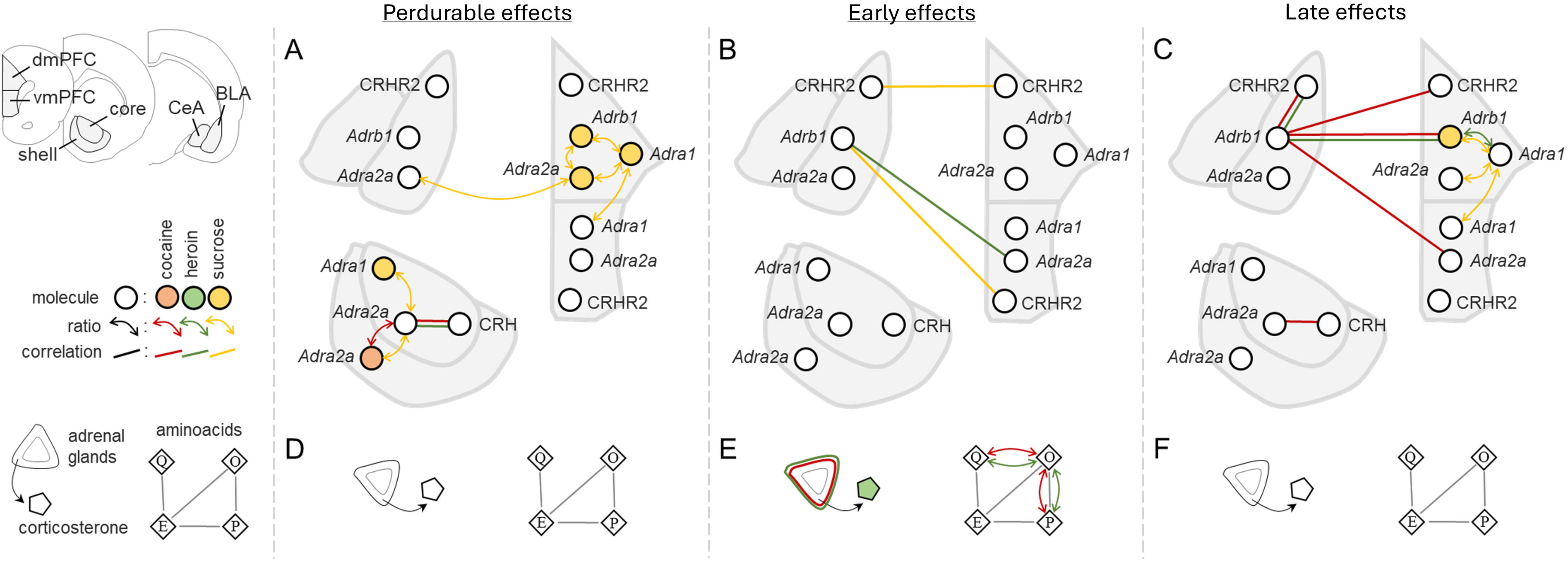
General scheme summarizing the perdurable (A, D), early (B, E), and late (C, F) effects in central (A-C) and peripheral (D-F) parameters after the incubation of cocaine (red), heroin (green), and sucrose (yellow) seeking. Individual molecules are depicted with a circle; ratios are depicted with a double headed arrow linking both parameters; correlations are depicted with a line connected both parameters. A grey line indicates a direct metabolic pathway between metabolites (Q: L-glutamine; E: L-glutamate; P: L-proline; O: L-ornithine). Abbreviations: dmPFC: dorsomedial prefrontal cortex; vmPFC: ventromedial prefrontal cortex; BLA: basolateral amygdala; CeA: central nucleus of the amygdala.

### 1. Adrenal hypertrophy at early drug withdrawal with concomitant increased corticosterone levels exclusively in heroin-exposed rats

In the present study, we observed a significant adrenal hypertrophy following cocaine and heroin—but not sucrose—self-administration, an effect typically associated with chronic stress (Ulrich-Lai and Herman, 2009). This alteration normalised after one month of withdrawal. Plasma corticosterone levels were transiently elevated after heroin self-administration, in line with previous reports in cocaine-exposed animals (Thiel et al., 2012). Neither we nor Grimm and colleagues (Grimm et al., 2016) detected comparable effects following sucrose self-administration (in their study, animals were anaesthetised prior to blood collection). Together, these findings suggest that, unlike sucrose, drug self-administration engages the hypothalamic–pituitary–adrenal (HPA) axis during early withdrawal under conditions that promote incubation of seeking (Fig. 6E). Notably, however, increased circulating corticosterone was observed only in heroin-exposed rats.

In humans, drug-related cues activate both the autonomic nervous system (ANS) and the HPA axis, with more pronounced responses in heavy users (Fox et al., 2005). Furthermore, both clinical and preclinical evidence indicates that drug exposure per se alters ANS and HPA axis function (Goldfarb and Sinha, 2018a; Ulrich-Lai and Herman, 2009). For instance, heroin intake suppresses HPA axis activity, whereas cocaine potentiates it, yet abstinence from either drug is associated with HPA axis hyperactivity (Brown et al., 2006). Consistently, rats with extended access to cocaine exhibit higher plasma corticosterone levels than those with restricted access (Mantsch et al., 2014). In human substance users, HPA axis dysregulation is frequently reported, characterised by elevated basal cortisol concentrations and exaggerated cortisol responses to stress (Goldfarb and Sinha, 2018b). By contrast, contingent consumption of natural reinforcers generally attenuates ANS and HPA axis activation, unless cues are presented in the absence of reinforcement, in which case both systems are recruited (Ulrich-Lai and Herman, 2009). Importantly, none of the aforementioned human studies on incubation examined whether cue-induced cardiovascular stress responses also exhibit incubation over time.

We also detected increased plasma L-ornithine to L-glutamine and L-ornithine to L-proline ratios, concomitant with adrenal hypertrophy following drug self-administration (Fig. 6E). This observation is of interest given evidence for a potential homeostatic interaction between glucocorticoids and L-ornithine metabolism. Oral L-ornithine administration reduces plasma glucocorticoid levels in stressed mice (Kurata et al., 2012) and in humans (Miyake et al., 2014), and corticosterone elevations in mice appear to be negatively regulated through L-ornithine metabolic pathways (Zamora-González et al., 2020). The increase in L-ornithine observed in the cocaine- and heroin-exposed groups may therefore reflect a compensatory response to HPA axis activation.

Additionally, further support for HPA axis engagement is provided by the endocannabinoid alterations observed in the same cohort of animals (Roura-Martínez et al., 2020). In this previous work, we reported persistent endocannabinoid-related molecular alterations detectable from the first day of abstinence and lasting for at least one month. Both cocaine and heroin reduced *Cnr1* expression in the dorsomedial prefrontal cortex. Cocaine additionally decreased *Napepld* and *Faah* expression in the nucleus accumbens shell, whereas heroin increased *Faah* expression in the ventromedial prefrontal cortex, consistent with diminished anandamide signalling. Interestingly, sucrose self-administration also induced molecular adaptations in a partially overlapping direction: increased *Dagla* and *Mgll* expression in the dorsomedial prefrontal cortex, reduced *Napepld* expression in the nucleus accumbens core, and increased *Faah* and *Cnr1* expression in the nucleus accumbens shell. Overall, this transcriptional profile is compatible with reduced anandamide tone and a relative enhancement of 2-AG signalling, a pattern reminiscent of chronic stress-induced endocannabinoid adaptations. Indeed, both chronic stress and exogenous corticosterone administration are known to induce region-specific changes in central endocannabinoid signalling (Bowles et al., 2012; Gray et al., 2016; Häring, Guggenhuber, & Lutz, 2012; McLaughlin & Gobbi, 2012; Ramikie & Patel, 2012), typically characterised by reduced anandamide tone and increased 2-arachidonoylglycerol (2-AG) levels. For instance, in the basolateral amygdala, these manipulations produce long-lasting decreases in anandamide signalling via FAAH up-regulation, together with transient elevations in 2-AG (Ramikie and Patel, 2012).

### 2. CRF and NA differential effects in the basolateral amygdala and the medial prefrontal cortex

Across all reinforcers, molecular changes converged after one month of withdrawal on *Adrb1* expression in both the basolateral amygdala (BLA) and the dorsomedial prefrontal cortex (dmPFC) (Fig. 6C). Both regions receive dense noradrenergic innervation from the locus coeruleus via the dorsal noradrenergic bundle; however, receptor dynamics and functional consequences differ markedly between them. ADRA2A, a high-affinity receptor preferentially engaged under conditions of low arousal, exerts excitatory effects in the prefrontal cortex but inhibitory effects in the BLA. Conversely, ADRB1, a lower-affinity receptor recruited during stress, is inhibitory in the prefrontal cortex but excitatory in the BLA (Arnsten, 2009; Buffalari and Grace, 2007; Ferry and McGaugh, 2008).

Evidence from reconsolidation paradigms further supports a functional role for β-adrenergic signalling within the BLA: pharmacological inactivation of β-receptors in this region disrupts reconsolidation of alcohol self-administration memories without affecting extinction processes (Chesworth and Corbit, 2018). The potential implication of ADRB1, rather than ADRA2A, in incubation is therefore of interest. In abstinent cocaine users, the late positive potential—a neurophysiological index of cue-induced craving that increases over abstinence—depends on β-adrenergic activation within the amygdala (De Rover et al., 2012; Parvaz et al., 2016), suggesting translational relevance for β-receptor-mediated mechanisms.

In the sucrose group, we observed a sustained shift in the balance of adrenergic receptor expression between the BLA and medial prefrontal cortex. Specifically, expression of all three adrenergic receptors was reduced in the dmPFC, with a proportionally greater reduction in the low-affinity receptor *Adrb1*. In parallel, the BLA displayed relatively higher *Adra2a* expression compared with the dmPFC, an effect that became more pronounced after one month of withdrawal. A comparable reduction in medial prefrontal adrenergic receptor expression has been reported in a mouse model of escalated ethanol consumption, accompanied by cognitive impairment during withdrawal, although causality was not established (Athanason et al., 2023).

We also detected a reduction in *Adra1* expression in the dmPFC relative to the ventromedial PFC after one month of withdrawal. Intra-medial prefrontal administration of the α1-adrenergic antagonist terazosin attenuates cocaine-primed reinstatement (Schmidt et al., 2017). However, given that the prelimbic cortex—targeted in that study—lies at the interface between dorsomedial and ventromedial subregions, and that their paradigm involved extinction followed by drug-primed reinstatement rather than incubation, direct comparisons are limited.

With regard to corticotropin-releasing hormone (CRH) signalling, we observed alterations in the relative balance between *Crhr2* expression in the BLA and mPFC and *Adrb1* expression in the BLA, suggesting coordinated adaptations within stress-related neuromodulatory systems. CRH neurons in the dorsal raphe projecting to the BLA have recently been shown to sustain cocaine memory strength following memory destabilisation (Ritchie et al., 2024). In a methamphetamine incubation model, *Crhr2* expression in the PFC was reduced after one month of abstinence in both sexes, with additional reductions in *Crhr1* in females; transcript levels were negatively correlated with lever pressing during withdrawal day 30 extinction testing (Daiwile et al., 2021).

Taken together, our findings suggest that both noradrenergic and CRH systems undergo region-specific adaptations that may alter their capacity to regulate substance seeking. These effects appear early in withdrawal in the sucrose group and become more evident during protracted withdrawal in cocaine- and heroin-exposed animals. This interpretation is consistent with extensive evidence implicating CRH in stress-induced reinstatement of heroin and cocaine seeking in rodents (Erb et al., 1998; Yavin Shaham et al., 1998, 1997). In alcohol-dependent individuals, CRHR1 antagonists have not demonstrated robust anti-craving effects during early withdrawal (Kwako and Koob, 2018; Schwandt et al., 2016b); however, their efficacy following prolonged abstinence remains largely unexplored.

### 3. Enduring effects on NA receptors in the nucleus accumbens

In the nucleus accumbens (NAc), we observed predominantly enduring alterations centred on *Adra2a* expression in the core subregion. Both cocaine and sucrose self-administration increased *Adra2a* expression in the core relative to the shell; in the sucrose group, this was accompanied by reduced *Adra1* expression. Following cocaine and heroin exposure, the correlation between *Adra2a* and *Crh* expression in the core was no longer present, suggesting a decoupling of noradrenergic and CRH-related transcriptional regulation in this region.

Preclinical literature has demonstrated that systemic or intracerebroventricular administration of α2-adrenergic agonists reduces cue-induced reinstatement of cocaine seeking (Smith and Aston-Jones, 2011), stress-induced—though not cocaine-primed—reinstatement of cocaine seeking (Suzanne Erb et al., 2000), and stress-induced heroin seeking (Yavin Shaham et al., 2000). Conversely, systemic administration of yohimbine, an α2-adrenergic antagonist that enhances noradrenaline release and induces anxiety-like responses, potentiates cocaine seeking when administered following extinction, albeit not in a classical incubation-like pattern (Shepard et al., 2004). As these manipulations were not region-specific, the precise contribution of α2 receptors within the NAc core remains unclear.

To our knowledge, the only study directly targeting adrenergic receptors within the NAc employed intra-accumbal administration of the α1-adrenergic antagonist terazosin, which did not alter cocaine-primed reinstatement (Schmidt et al., 2017). Moreover, cocaine-primed reinstatement was likewise unaffected by α2 agonist administration. In clinical populations, however, α2-adrenergic agonists reduce both stress-and cue-induced craving in opioid- and cocaine-dependent individuals (Hermes et al., 2019; Jobes et al., 2011b; Kowalczyk et al., 2015b; Sinha et al., 2007a), highlighting a potential translational dissociation between systemic and region-specific mechanisms.

### 4. A proposed mechanistic framework

Taken together, the present findings support a model in which withdrawal from drug self-administration engages stress-related neuromodulatory systems that progressively reshape corticolimbic circuitry implicated in cue-driven seeking. Early withdrawal was characterised by adrenal hypertrophy and transient corticosterone elevation (in heroin-exposed rats), consistent with activation of the hypothalamic–pituitary–adrenal (HPA) axis under conditions known to promote incubation of seeking. These peripheral adaptations were paralleled by central transcriptional changes within the corticotropin-releasing hormone (CRH) and noradrenergic (NA) systems.

At the cortical–amygdalar level, convergent alterations in *Adrb1* expression after one month of withdrawal point to a sustained reconfiguration of β-adrenergic signalling in both the basolateral amygdala (BLA) and dorsomedial prefrontal cortex (dmPFC). Given the opposing functional effects of β1 receptor activation in these regions—facilitatory within the BLA and inhibitory within the prefrontal cortex— (Arnsten, 2009; Buffalari and Grace, 2007; Ferry and McGaugh, 2008) such adaptations may bias network dynamics towards enhanced emotional salience (Kong et al., 2025) and diminished top–down control under stress. In parallel, region-specific changes in *Crhr2* expression and altered correlations between CRH- and NA-related transcripts suggest a reorganisation of stress-responsive signalling rather than isolated receptor regulation.

Within the nucleus accumbens (NAc), enduring increases in *Adra2a* expression in the core subregion, together with the loss of correlation between *Adra2a* and *Crh* after cocaine and heroin exposure, indicate a potential decoupling of noradrenergic and CRH modulation at a key integrative node of motivated behaviour. Given the established role of the NAc core in cue-driven reinstatement and action selection (Wang et al., 2010), these molecular changes may influence the integration of stress and reward signals during protracted withdrawal.

Importantly, although sucrose self-administration did not induce overt peripheral markers of HPA activation, it produced partially overlapping central adaptations, particularly at early withdrawal stages. This suggests that repeated reward-driven behavioural engagement may recruit stress-related neuromodulatory systems at the circuit level even in the absence of systemic endocrine activation. In contrast, drug self-administration was associated with more pronounced or persistent alterations during protracted withdrawal, consistent with the notion that drugs of abuse co-opt stress circuitry to a greater extent than natural reinforcers.

Collectively, these findings support a framework in which withdrawal-related stress signalling progressively shifts the balance between prefrontal regulatory control, amygdalar emotional encoding, and accumbal action selection (Balleine and Ostlund, 2007). Such circuit-level reorganisation may provide a mechanistic substrate for the time-dependent intensification of cue-driven seeking.

### 5. Limitations and suggested new research

The present study was conducted exclusively in male rats, which represents a limitation. Female rodents often display enhanced stress responsiveness and differential CRF–noradrenergic coupling (Bangasser et al., 2019; Bangasser and Wiersielis, 2018), factors that may critically influence the incubation of reward seeking. Future studies should determine whether the molecular adaptations described here generalise to females and whether sex hormones modulate these withdrawal-associated transcriptional signatures. A second aspect to consider as a potential limitation is the differences between the sucrose and drug self-administration groups (e.g., session length, route of administration, and non-equivalent control conditions). Even if these differences reflect the specific nature of each type of reinforcer of the published protocols to induce incubation, they introduce potential confounds that must be taken into consideration. A third caveat, which is a common criticism in incubation studies, concerns whether observed molecular adaptations are causally related to the progressive increase in cue-induced responding, or merely epiphenomenal correlates of withdrawal. Several aspects of the present data strengthen their relevance to the incubation phenomenon (Roura-Martínez et al., 2020). First, key adaptations emerged or converged during protracted withdrawal, temporally aligning with the period in which incubation has emerged or has even reached its peak (Grimm et al., 2001). Second, affected regions—BLA, dmPFC, and NAc core—are core components of the circuitry mediating cue-induced reinstatement and incubation across substances (Chow et al., 2025). Third, the receptors most consistently altered (β1- and α2-adrenergic receptors, as well as CRH receptors) are known modulators of stress-induced and cue-induced reinstatement in both rodent models and clinical populations (see Introduction).

Nevertheless, the present study is purely descriptive and does not establish functional causality. Future experiments employing region-specific pharmacological or chemogenetic manipulations during defined withdrawal time points will be required to determine whether the identified adaptations are necessary or sufficient for the incubation phenomenon. In particular, selectively modulating β1-adrenergic or CRH receptor signalling within the BLA–mPFC– NAc network during protracted withdrawal would allow direct testing of the proposed model.

## ACKNOWLEDGEMENTS

We would like to thank Rosa Ferrado for the excellent technical assistance.

## CONTRIBUTORS

D.R-M performed the experiments, the statistical analyses and wrote the first draft of the manuscript.

M.U. contributed to the statistical analysis and the first draft of the manuscript.

M. M-F, C. A C. and I B-Y performed part of the molecular experiments.

A.B. performed the capillary electrophoresis analysis.

E. A. contributed to secure funding and oversaw the project

Alejandro Higuera-Matas designed the experiments, secured funding and wrote the final version of the manuscript.

**Tables 1 and 2**. Physiological, biochemical, and neuromolecular profiles during withdrawal in rats exposed to saline, cocaine, or heroin (**Table 1**) and sucrose or water (**Table 2**). Data represent descriptive and inferential statistics for body and adrenal weights, plasma markers (corticosterone and amino acids), and mRNA expression of stress-related genes (e.g., *Crh*, *Crhr2*) and adrenergic receptors (*Adra1*, *Adra2a*, *Adrb1*), including their ratios, as well as protein expression for specific targets (CRH and CRHR2), across distinct brain regions: nucleus accumbens (shell and core), prefrontal cortex (dmPFC and vmPFC), and amygdala (BLA and Central Amygdala -CeA). Rats were evaluated at two time points: 1 day (wd1) and 30 days (wd30) of withdrawal. Descriptive statistics are presented as mean ± standard deviation (SD) for n = 6-8 per group. Inferential analysis was performed using a two-way ANOVA (drug x withdrawal period), with main effects and interactions reported as F-statistics, p-values, and partial eta-squared (η²p) as a measure of effect size. Post-hoc (REGWQ) comparisons were conducted to identify significant differences between treatment groups (saline, cocaine, and heroin). Interactions were analysed by simple effects analysis with Sidak correction. An ‘x’ indicates missing data or data that did not comply with ANOVA assumptions after all possible data transformations. Statistical significance was set at p < 0.05.

**Table 3**. Correlation analysis and statistical comparison of gene expression relationships across brain regions. The table displays Pearson correlation coefficients (r) between mRNA expression or protein levels of stress-related genes and adrenergic receptors in specific brain regions (e.g., *Crhr2*, CHR2, CRH, *Adrb1*, *Adra2a* within the dmPFC, vmPFC, BLA, and NAcc core). Data are categorized by experimental group (Water, Sucrose, Saline, Cocaine, and Heroin) and withdrawal period (1 day [wd1] and 30 days [wd30]). For each correlation, the coefficient (r), p-value (p), and sample size (N) are provided. Statistical comparisons between correlation coefficients were performed using Fisher’s r-to-Z transformation. The resulting Z-scores (Z) and corresponding p-values (p) indicate the significance of differences between withdrawal periods within the same treatment (e.g., wd1 vs. wd30) and between treatment groups at a specific time point (e.g., saline vs. cocaine). Statistical significance for both correlations and Z-tests was defined as p < 0.05.

